# Physiological Artefacts and the Implications for Brain-Machine-Interface Design

**DOI:** 10.1101/2020.05.22.111609

**Authors:** Majid Memarian Sorkhabi, Moaad Benjaber, Peter Brown, Timothy Denison

## Abstract

The accurate measurement of brain activity by Brain-Machine-Interfaces (BMI) and closed-loop Deep Brain Stimulators (DBS) is one of the most important steps in communicating between the brain and subsequent processing blocks. In conventional chest-mounted systems, frequently used in DBS, a significant amount of artifact can be induced in the sensing interface, often as a common-mode signal applied between the case and the sensing electrodes. Attenuating this common-mode signal can be a serious challenge in these systems due to finite commonmode-rejection-ratio (CMRR) capability in the interface. Emerging BMI and DBS devices are being developed which can mount on the skull. Mounting the system on the cranial region can potentially suppress these induced physiological signals by limiting the artifact amplitude. In this study, we model the effect of artifacts by focusing on cardiac activity, using a current-source dipole model in a torso-shaped volume conductor. Performing finite element simulation with the different DBS architectures, we estimate the ECG common mode artifacts for several device architectures. Using this model helps define the overall requirements for the total system CMRR to maintain resolution of brain activity. The results of the simulations estimate that the cardiac artifacts for skull-mounted systems will have a significantly lower effect than non-cranial systems that include the pectoral region. It is expected that with a pectoral mounted device, a minimum of 60-80 dB CMRR is required to suppress the ECG artifact, while in cranially-mounted devices, a 20 dB CMRR is sufficient, in the worst-case scenario. The methods used for estimating cardiac artifacts can be extended to other sources such as motion/muscle sources. The susceptibility of the device to artifacts has significant implications for the practical translation of closed-loop DBS and BMI, including the choice of biomarkers and the design requirements for insulators and lead systems.

## I. INTRODUCTION

Deep brain stimulation (DBS) has been proven to be an effective therapy for neurological disorders such as Parkinson’s disease, essential tremor and Epilepsy. Furthermore, closed loop deep brain stimulation based on sensing local field potential signals (LFP) has shown the potential to deliver patient and state specific stimulation with improved results, power consumption and reduced side effects [1], [2].

DBS and brain-machine-interfaces (BMI) have two common placements, which are: 1) Pectoral mounted devices: these are devices implanted in the chest under the skin and below the collarbone, with electrodes tunneled through the neck to the area of interest in the brain. This is the currently the most common placement for DBS. 2) Cranial mounted devices: these devices are implanted on or in the skull under the skin with electrodes fed through to the area of interest in the brain. This approach is the less common of the two, but is used in the RNS system by Neuropace [3], and is being explored for emergent brain stimulators for research [4]. Fig. 1 illustrates the two different BMI placements.

**Fig. 1.**
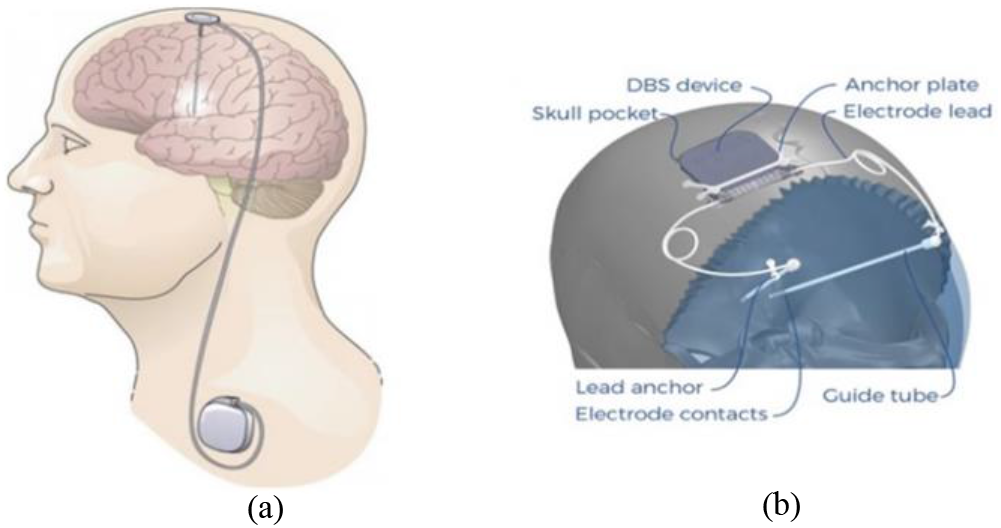
DBS Brain-Machine-Interface placements. (a) Pectoral mounted DBS device with electrodes extensions through the neck to the area of interest in the brain. (b) Cranial mounted DBS device. Source: Adapted from [4], [9].

In BMIs and closed-loop DBS stimulation, the precision sensing of the LFP or ECoG signals is essential to detect and extract the required biomarkers. However, recording these signals can be a difficult task due to persistent artifacts such as stimulation artifacts, electrocardiography (ECG) artifacts, and muscle movement artifacts [5]. These artifacts can mask the LFP signal and the underlying biomarkers. Most of the research in closed loop DBS is focused on removing and suppressing stimulation artifacts, which is prominent in all closed-loop DBS devices and bi-directional BMIs [6], [7], [8]. This issue is easily observed on the bench. An overlooked issue is ECG and muscle movement artifact, which requires a suitable model for testing, and is becoming more prominent as sensing systems are deployed commercially.

## II. MEASUREMENT BACKGROUND

Local field potentials are usually measured as a differential signal using the same DBS electrodes as for stimulation. The LFP signal sensed with a DBS electrode can range from 1-20 μVrms [10]. Most LFP oscillations are in low frequency bands, ranging from 2 Hz to 100 Hz, but they may go as high as 350 Hz [1].

When a DBS device is implanted in the chest, the device case can act as the system’s reference. The case is in a close proximity to the heart, which acts as a large signal generator that is superimposed on top of the brain signals of interest. The relatively large ECG signals are in the range of 0.5 mV to 5 mV with a frequency range of 0.05-150 Hz [11], [12]. These ECG artifacts are three orders of magnitude larger than the LFP signals of interest, with a frequency content that overlaps with the LFP frequency range.

The ECG can be modelled as coupling into the sensing input chain as a common mode signal. Fig. 2 illustrates an example of a differential LFP sensing circuit used in a DBS device. To suppress ECG artifacts in a chest implanted device, a front-end amplifier with a high common mode rejection ratio (CMRR) is required. However, it is quite challenging to achieve a high CMRR in an implantable system, even with commercially available high-performance instrumentation amplifiers (INA) with a CMRR larger than 100 dB. The CMRR can be undermined by the input impedance mismatch between the tissue-electrode interface and the front-end amplifier. The impedance mismatch can be caused by 1) impedance mismatch in the electrode tissue interface, 2) impedance mismatch along the lead and extension, 3) mismatch in DC blocking capacitors/ input high-pass filter (a common building block in DBS device), and 4) impedance mismatch in front-end passive low-pass filter components. In practice, shunt variations are the most likely issues to impact the CMRR in practice.

**Fig. 2.**
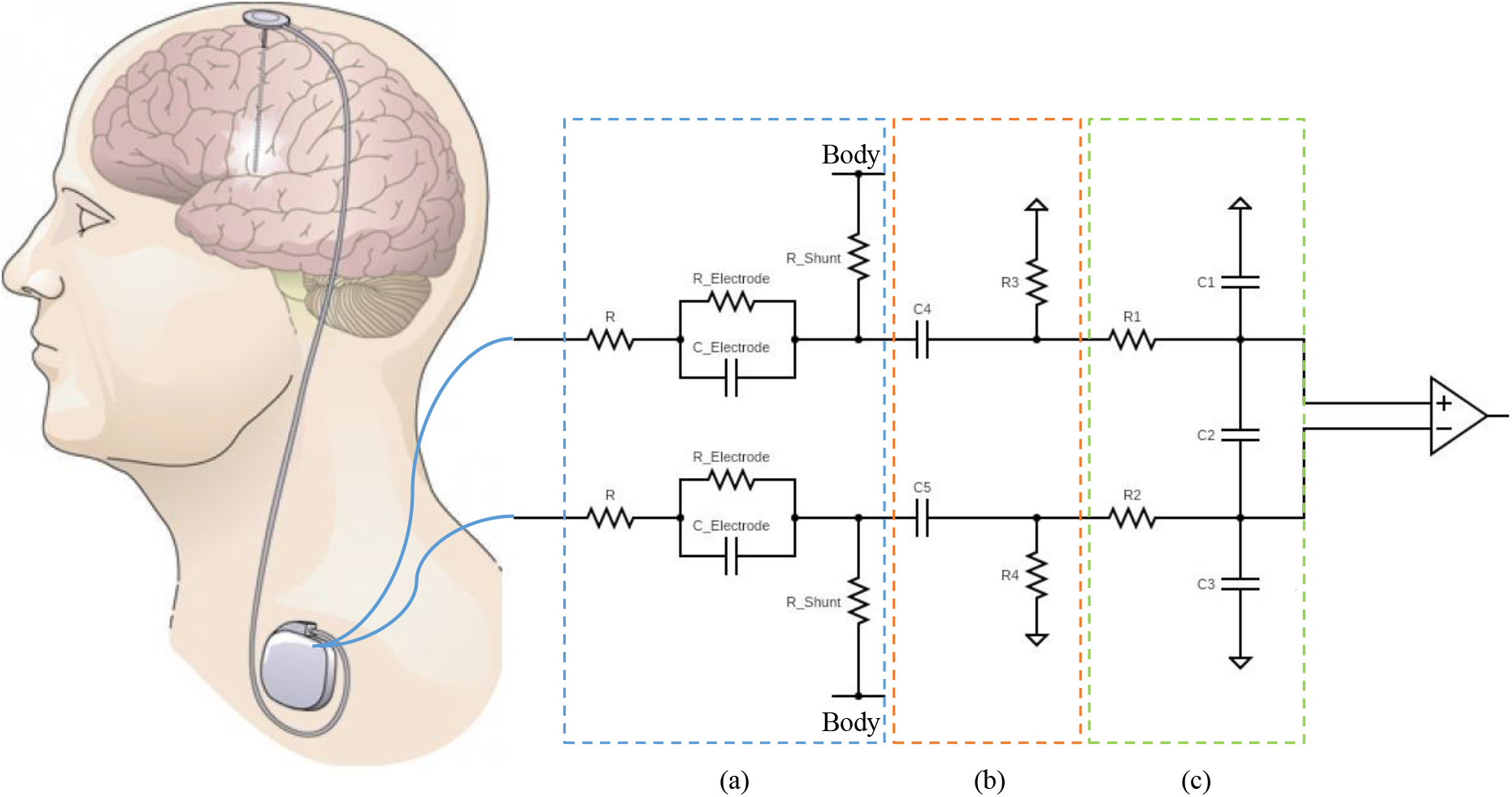
An example of a LFP differential sensing circuit used in DBS devices (a) Tissue electrode interface equivalent circuit and lead/extension routing, (b) DC blocking/high-pass filter, (c) passive low-pass differential filter. Source: Adapted from [9].

This paper looks to explore the relative impact of chest versus skull mounted DBS and BMI placements on sensing sensitivity to cardiac artifacts. By modelling the heart as a dipole source of the ECG artifact, the effect of the device placement on the induced current density, electric field and potential can be investigated. Using relative comparisons, a criterion for designing a differential amplifier will be presented for guiding the design of BMI and DBS systems.

## III. MODELING METHODS

### A. The Dipole model for cardiac activity in a uniform medium

The hypothetical model for cardiac electrophysiology underlying almost all the clinical devices for recording heart activity or ECG is that the heart is treated as a single current dipole source (which is equivalent to multiple dipole generators) and that the thorax has a uniform electrical conductivity [13]. Despite its simplicity, the equivalent dipolar source can be used in modelling ECG potentials and myocardial alterations [14].

### B. Problem statement

The conceptual idea is that cardiac activity can be modelled by a current-dipole source of variable orientation at a fixed location 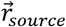. For an infinite isotropic and homogeneous medium with conductivity *σ_m_*, the current *I_source_* is created by the cardiac source located in the heart region Ω_*source*_ (the myocardium). The problem is to estimate the induced extracellular electric field 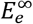 at some point external to such a dipole surface 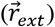, at pre-defined distances 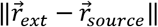. The objective is to then derive a formula to compute an induced artifacts as a function of the dipole and the spatial distance.

The induced electric field magnitude *E∞,* which is proportional to the extracellular potential and induced ECG artifact in the homogeneous medium [15], [16], is given by a volume integral over the sources:

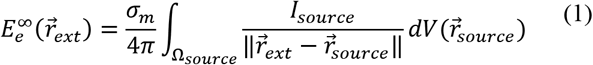

where the volume current density of the dipole is defined as 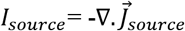. By applying the divergence

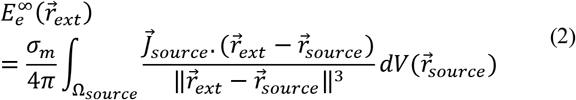

where the current density is zero on the boundary of the heart region. For numerical computations, the domain Ω_*source*_ is segmented into m sub-domains Ω_*n*_,*n* = 1,2, …,*m* (called elements), and (2) can be discretized as

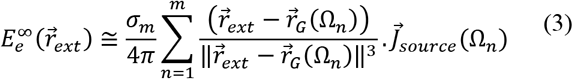

where 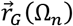 is the center of gravity of the sub-domain Ω_*n*_, and 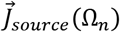 is

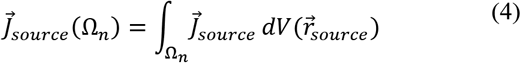

Equation (3) depicts the E-field as a superposition of the dipolar fields. Therefore, it is assumed that the equivalent heart dipole can rotate in space and generate three independent current densities (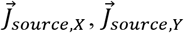 and 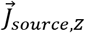) during the cardiac cycle. The superposition of these rotations is selected as the effective value of the artifacts that would capture any effect, which is valid for any linear medium.

### C. Torso model and solver

In this research, a three-dimensional MRI-derived torso model is utilized. The approximated heart model is defined by a homogeneous 7 cm diameter sphere consisting of 2 mm^3^ elements. The location of the heart is chosen according to the human anatomy and is surrounded by a homogeneous volume conductor (electrical conductivity = 033 S/m) with the shape of the human torso. The model was solved using a linear solver on COMSOL with finite element methods (FEM) using a relative error of 1.0e-6. Conventionally, the homogeneous conducting medium is introduced as a rational approximation to the electrical behaviour of human body tissues [17], [18]. The overall design framework of the proposed model of the heart and torso is shown in Fig. 3. The hypothetical locations of the dipole points are examined in two different scenarios: in the first scenario (I) the dipolar location is close to the chest and in the second scenario (II) close to the shoulder, as expressed in Fig. 3(b). According to (3), mesh cells are generated by a subdivision of a continuous geometric space into discrete geometric cells (Ω_*n*_), as illustrated in Fig. 3(b). For all simulations, the magnitude of the electric current dipole moment is assumed to be 1 (mA. meter), as suggested in [19], for the maximum possible value.

**Fig. 3.**
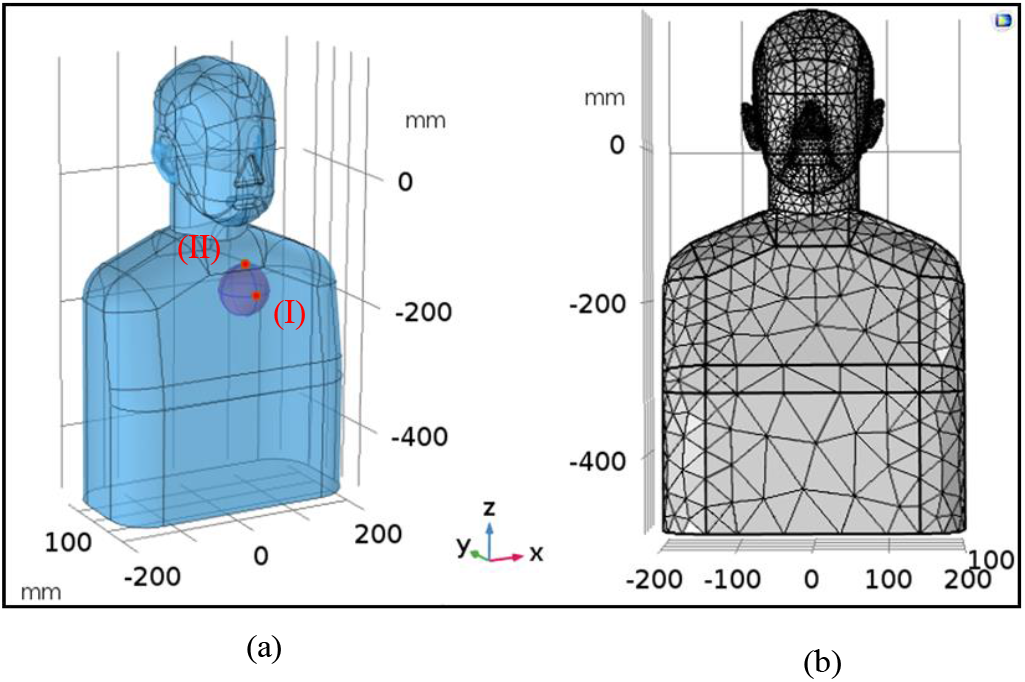
3D torso and heart model. (a) Torso and heart geometry used for the FEM. The purple sphere inside the chest indicates the heart model and the red spots denote the hypothetical locations of the current dipole. (b) The finite element mesh used to subdivide the torso model into m sub-domains (Ω_*n*_, *n* = 1,2, …,*m*).

## IV. RESULTS

### A. The activation map

The induced current and electric field during the cardiac cycle in the pectoral region, as a map of dipole activity in the first scenario, is shown in Fig 4. According to the FEM simulation results, the maximum induced current and electric field on the chest for oscillations in three directions are equal to 5.02 A/m^2^ and 15.2 V/m, respectively.

**Fig 4.**
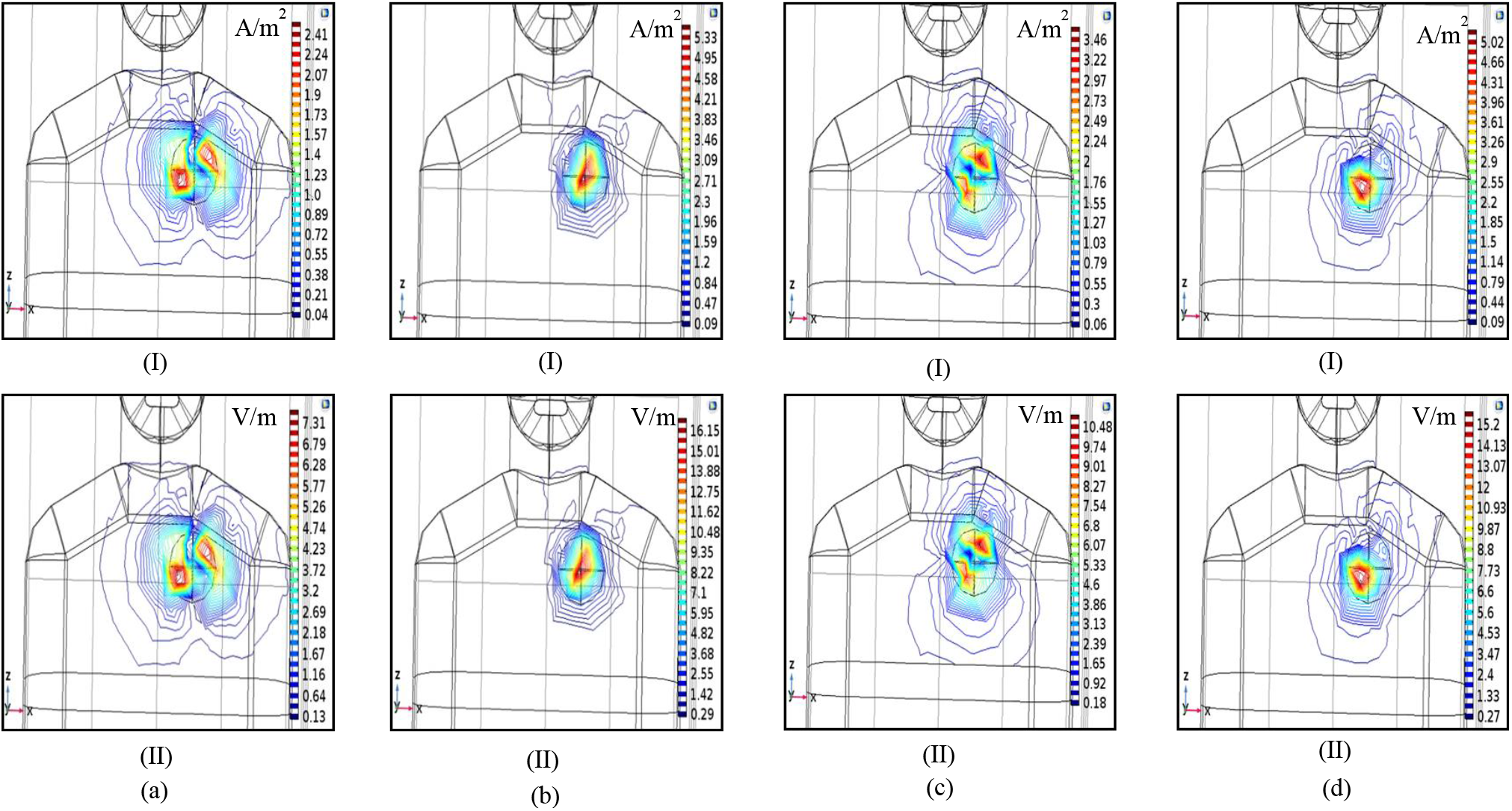
Cardiac activation maps in the pectoral region. Induced current density (I) and electric field (II) for dipole oscillations: (a) Only in the X direction 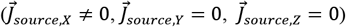. (b) only in the Y dilution 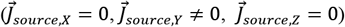. (c) only in the z dilution 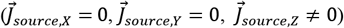. (d) In all directions, as a superposition.

Fig. 5 displays the induced current and E-field as induced ECG artifacts at the skull level. Evidently, the maximum induced artifacts are seen in the temporal region, which is due to the proximity to the neck and the anatomical location of the heart. It is noteworthy that in these figures, the values of the induced artifacts on the viscerocranium (or facial skeleton) are not shown.

**Fig. 5.**
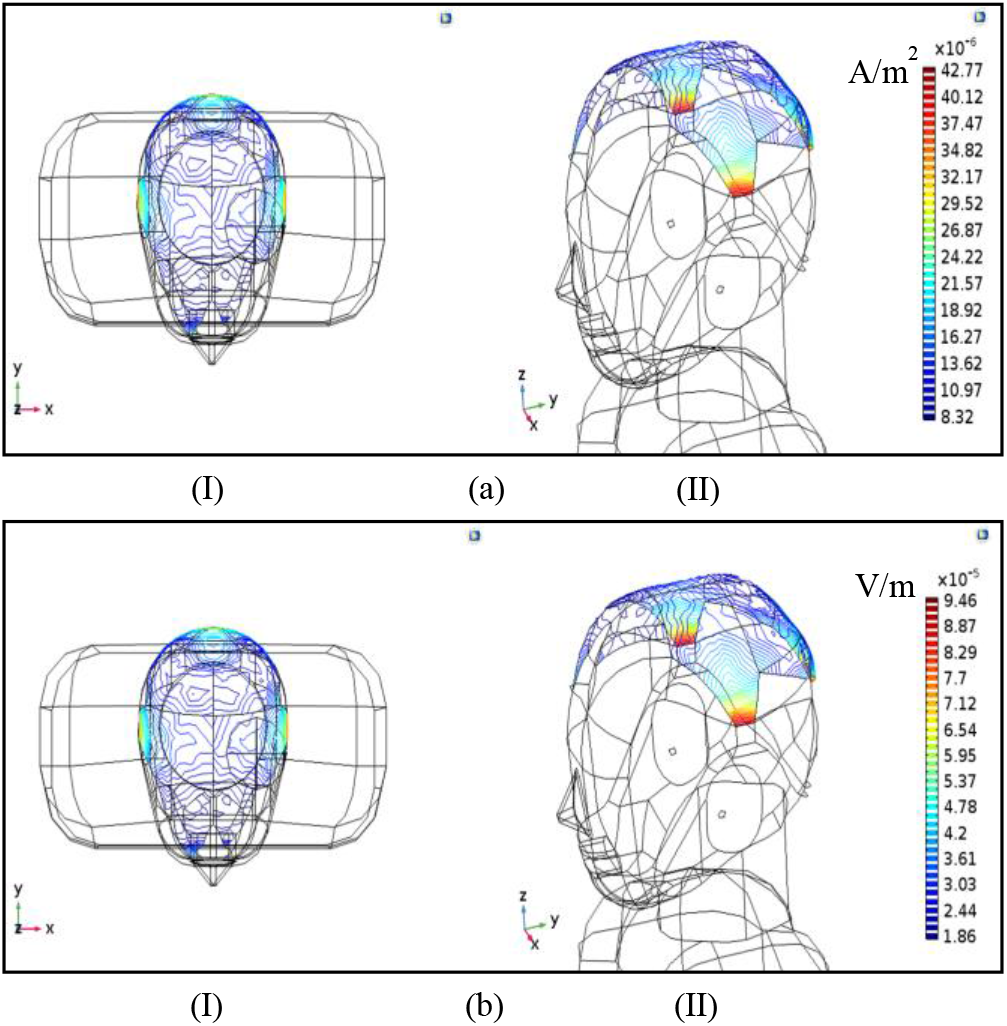
ECG artifact at the skull. Induced current density (a) and electric field (b) for dipole oscillations in all directions 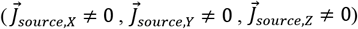, as a superposition.

As shown in Fig. 5, the maximum induced current and electric field at the skull for the superposition state are approximately equal to 42.7 μA/m^2^ and 94 μV/m, respectively. Compared to the artifact values around the heart, these parameters have been significantly attenuated. As a criterion of the induced artifact at the skull, the two rates are defined by the following formulas.

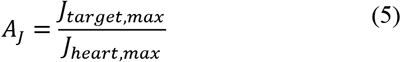

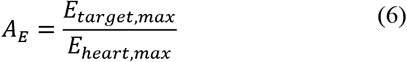

where A_j_ and A_E_ are the induced current density and the E-field artifacts rate, J_heart,max_ and E_heart,max_ are the maximum current density and the maximum E-field around the heart, and J_target,max_ and E_target,max_ are the maximum induced values at the target region, respectively. According to Fig. 3(a), the dipole locus can be assumed in two different scenarios and then the values of the artifacts can be calculated. The results of these two scenarios and artifact values in different areas of the skull are described in Fig. 6.

**Fig. 6.**
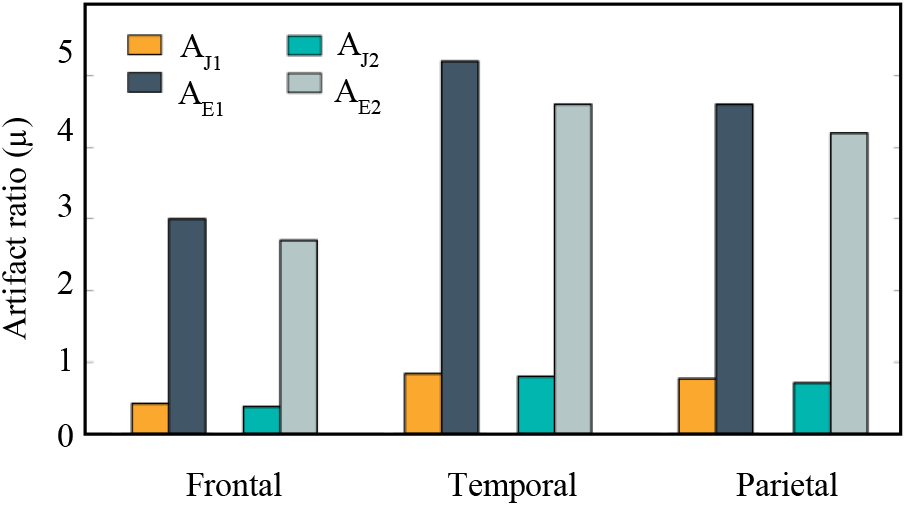
Estimated artifact ratios in different areas of the skull; includes Frontal, Temporal and Parietal regions, where AJ1 and AJ2 denote the current density artifact ratios for the first and second scenarios, AE1 and AE2 indicate the E-field artifacts for the first and second scenarios, respectively.

The values in Fig. 6 depict how the ratios of the artifacts in the cranial area change as the current-dipole source location changes between two scenarios. But in the worst-case scenario, artifact ratios are expected to decay significantly (1:0.5μ).

As a criterion for designing a differential amplifier and necessary CMRR, the potential difference values of cardiac activity have been simulated using the path integral for DBS devices located in the chest and the skull, as shown in Fig. 7. These values indicate how much of the common-mode ECG artifact will appear in each DBS system. According to the simulation results, in the worst-case scenario, the ratio of the common-mode ECG signal in the chest-mounted DBS system is 2700 times that of the skull-mountable DBS device (V2/V4).

**Fig. 7.**
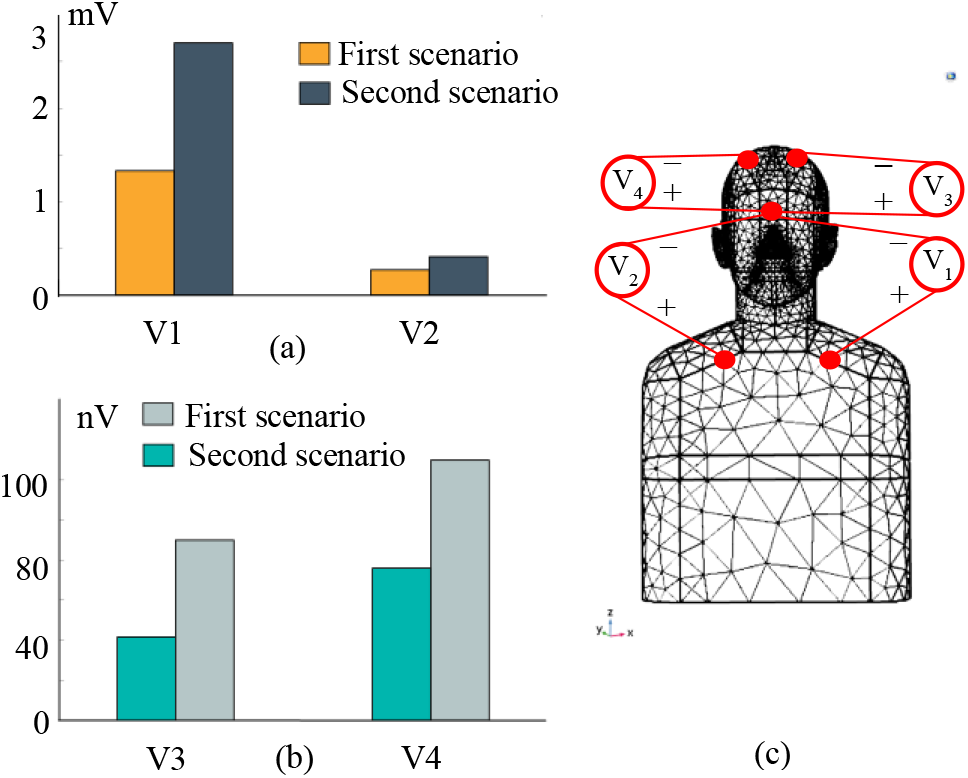
The estimated potential difference for the first and second scenarios. (a) Chest cavity to a distant point in the middle of the brain (V1 and V2), as a common-mode ECG artifact for a chest mounted DBS and BMI systems, as shown in Fig. 1a. (b) From the same centre point to the top of the head (V3 and V4), as a metric used to quantify common mode ECG artifact for a cranial-mount DBS and BMI systems, as shown in Fig. 1b. (c) Measuring locations of potential differences in the FEM model. V1 and V2 measurements were performed at a depth of 3 cm inside the pectoral region (Y direction). V3 and V4 potential differences were executed at a depth of 4 cm inside the head (Y direction).

## V. MODEL INTERPRETATION

The large reduction in common mode ECG artifacts with the two different placement results in a reduction of the required system CMRR. Recall that typical field potential measurements are on the order of 1-20 μV⊓⊓s. In a pectoral-mounted device, a 60-80 dB CMRR is therefore required to supress these levels of ECG artifacts. On the other hand, in a cranial mounted DBS device, a 20 dB CMRR is sufficient to supress these artifacts in the worst-case scenario. To help provide context for these numbers, a typical platinum-iridium electrode has an impedance of approximately 1θ k-100 kΩ in the low frequency bands of interest. Using the model in Fig. 2, we can model the CMRR as the mismatch of electrodes along the pathway.

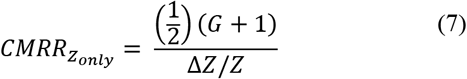

where CMRR_Zonly_ is the CMRR due only to tissue electrode interface and shunt impedance mismatch (ideal Amplifier case), *△%/%* Is the impedance matching ratio and G is the nominal ratio Z_tissue_electrode_/Z_shunt_. [20]; in general, this should trend to zero in a well-insulated lead and extension.

To reject these far-field artifacts, we must maximize the CMRR. As a design heuristic, to maintain a 60 dB CMRR, the shunt impedances along the lead and extension must therefore match to within 10-100 MΩ. This level of isolation is challenging for polymer-coated electrodes chronically exposed to the saline environment. To further maximize the equivalent shunt resistance, we would ideally float the pulse generator case to maximize its isolation. In practice, this can be challenging in brain stimulation systems that use “monopolar” stimulation between the electrode and the case. The best-case scenario is then to use active recharge, which limits the time duration of the case connection [21]. One opportunity to relax these requirements is to lower the tissue-electrode impedance with surface coatings like PEDOT or Ti-N [22], [23]. On the other hand, the cranial system requires only 200 k-2 MΩ, which is much more in line with standard industry capability and requires no new material interfaces for acceptable artifact rejection.

Additional mitigations are to explore the use of additional sealing adhesives at the joints between leads, extensions and the device connector block. While these might provide improved isolation, the additional surgical complexity and impact on revisions/replacements needs to be carefully considered.

Another opportunity for lowering susceptibility to artifacts is to explore alternative biomarkers. One interesting candidate is the evoked potential, which is the response of the neural circuit to stimulation and which can have an amplitude in the common DBS targets of several 100 microvolts. Recent research suggests that such evoked potentials can encode information about the location of the electrode and the brain’s state [24]. The advantage of these signals is that they use methods of synchronous detection to avoid artifacts. Similar to chopper methods to remove low-frequency offset and drift [25], if the stimulation frequency is above the cardiac energy, then the impact of the cardiac artifact is greatly attenuated. The major trade-off of this technique is the bandwidth required for sampling, typically ten times higher than for low frequency field potentials, and the need to manage stimulation artifacts to allow for resolution of physiological signals within milliseconds of stimulation termination [24], [26]. However, sampling rates are well within the control of the design engineer and fall within typical rates found commonly in implant systems [27].

## VI. DISCUSSION

There are several considerations in choosing between architectures, and each design has limitations and trade-offs. The cranial mounted systems are limited in terms of implant location on the skull, which in turn limits the size of the device. This limitation usually results in a compromise in battery size, which affects the battery life of the device. However, battery size and life in implantable devices are becoming less of an issue, since the approval of rechargeable DBS systems, although recharge frequency might be impacted.

Models such as this one might also suggest optimal surgical pathways to use for the system. For example, the model suggests that the right-side pectoral region might be a more optimal location to place a sensing device. On the cranium, it has been shown that the Frontal area will have the least value of ECG artifacts. Therefore, this region is recommended as an acceptable cranial area for DBS and BMI devices mounting, in terms of minimum induced ECG artefact, if surgical placement is feasible.

One issue to highlight is that the impedances that impact the CMRR and artefact susceptibility are orders of magnitude larger than the tissue-electrode impedance. This means that sensing path mismatch will often be difficult to diagnose with the relatively crude impedance checks performed with existing medical implants.

Finally, it is worth noting that the dipole model can be used for multiple artifacts including muscle signals. In general, the same principles will apply, and the reduced muscle content of the cranium will aid in lowering artifact sensitivity.

## VII. LIMITATIONS

This work also has several limitations, which must be taken into consideration. For instance, a simplified body model is utilized while human body tissues have inhomogeneous anisotropic conductivity. Examining and selecting inhomogeneous materials for the torso and considering the possibility of transmitting cardiac signals with blood vessels, as a model closer to the real body medium, may increase the value of artifacts seen on the skull. Defining biologically-based brain models with different layers (including skull, scalp, cerebrospinal fluid, grey matter, and white matter) [28], can better represent the propagation model of artifacts on the cranial region and the DBS lead.

Due to the complex geometry and different boundary conditions in body materials with separate electrical parameters, it is very complicated to find an analytical solution. The FEM can provide approximate answers to analytical solutions over the predefined geometry. Increasing the mesh density can reduce computational error, although it increases overall analysis time. Another important consideration is how to model the cardiac activity. The present work uses the current-source dipole model and applies the rotated current dipole moment as a cardiac cycle, but there are other models introduced in [16], [29], which can generate different parts of the ECG signal (so-called QRS complex) and represent ventricular depolarization. These models enable a conceptual basis for a deeper understanding of the ECG artifacts in the pectoral and the cranial regions. Therefore, in future studies, by considering these cases, a more accurate estimate of the ECG artifacts can be calculated.

## VIII. SUMMARY

In this study, we show the importance of placement of DBS devices and BCI where LFP signal sensitivity is paramount. As illustrated by the numerical modelling and finite element method results, cranial mounted DBS-BMI devices are less susceptible to ECG artifacts, which in turn reduces the CMRR system requirements for the implantable device. This makes cranial mounted systems a preferable choice for recording low frequency physiological signals without interference from ECG artifacts. However, installing DBS-BMI devices in the cranial area may require a smaller system, resulting in lower battery life or increased recharging burden for the patient. Paying attention to lead and extension design, as well as specific placement in the torso, can help improve the probability of artifact free sensing with a pectoral implant. Finally, the choice of biomarker might also prove critical. While low frequency field potentials overlap with ECG, the application of evoked potentials might provide the same sensitivity advantages as chopper stabilization methods and provide artifact-free measurements. Ultimately, the balance of clinical validity and technical feasibility will determine translational success of Brain-Machine-Interfaces.

